# Quantitative super-resolution single molecule microscopy dataset of YFP-tagged growth factor receptors

**DOI:** 10.1101/246488

**Authors:** Tomáš Lukeš, Jakub Pospíšil, Karel Fliegel, Theo Lasser, Guy M. Hagen

## Abstract

**Background:** Super-resolution single molecule localization microscopy (SMLM) is a method for achieving resolution beyond the classical limit in optical microscopes (approx. 200 nm laterally). Yellow fluorescent protein (YFP) has been used for super-resolution single molecule localization microscopy, but less frequently than other fluorescent probes. Working with YFP in SMLM is a challenge because a lower number of photons are emitted per molecule compared to organic dyes which are more commonly used. Publically available experimental data can facilitate development of new data analysis algorithms.

**Findings:** Four complete, freely available single molecule super-resolution microscopy datasets on YFP-tagged growth factor receptors expressed in a human cell line are presented including both raw and analyzed data. We report methods for sample preparation, for data acquisition, and for data analysis, as well as examples of the acquired images. We also analyzed the SMLM data sets using a different method: super-resolution optical fluctuation imaging (SOFI). The two modes of analysis offer complementary information about the sample. A fifth single molecule super-resolution microscopy dataset acquired with the dye Alexa 532 is included for comparison purposes.

**Conclusion:** This dataset has potential for extensive reuse. Complete raw data from SMLM experiments has typically not been published. The YFP data exhibits low signal to noise ratios, making data analysis a challenge. These data sets will be useful to investigators developing their own algorithms for SMLM, SOFI, and related methods. The data will also be useful for researchers investigating growth factor receptors such as ErbB3.

## Context

Fluorescence optical microscopy is one of the most important tools available for the study of biological systems at the cellular level. Unfortunately, due to diffraction phenomena the resolution of fluorescence microscopes in the lateral *d* dimension is limited to

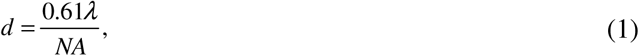

where λ is the wavelength of the detected light, and NA is the numerical aperture of the objective lens. As many biological structures within cells are much smaller than this, increasing resolution is of prime importance. Today several methods have been developed which are able to image below the diffraction limit [1,2].

Photoactivated localization microscopy (PALM) [3] was initially accomplished with the photoconvertible fluorescent protein mEOS [4]. A similar method, (direct) stochastic optical reconstruction microscopy (d)STORM utilizes organic dyes [5–8]. In these super-resolution methods, single fluorescent molecules are induced to blink on and off (photoswitching) randomly in the sample. A sensitive camera is used to record an image sequence of the single molecule blinking events, and a computational algorithm is used to fit the imaged point spread functions (PSFs) to a model function [9,10]. By doing so, the coordinates of each molecule can be determined with an uncertainty which is below the diffraction limit [11]. Once enough molecules have been imaged (usually 10^6^−10^7^ are required, depending on the sample structure [12]), an image can be reconstructed with lateral resolution improved by about a factor of 10. This is done by plotting the coordinates of each molecule in a new image with a much smaller pixel size. Together, this family of methods is known as single molecule localization microscopy (SMLM).

Although PALM experiments were initially performed with fluorescent proteins which are specifically photoconvertible [3], green fluorescent protein (GFP) and its spectral variant yellow fluorescent protein (YFP) are also known to exhibit blinking characteristics [13]. GFP and YFP have been used in SMLM, but less frequently [14–20]. Here we used a modified YFP known as mCitrine [21] for SMLM. The advantage of using mCitrine is that SMLM can be accomplished with a single laser, rather than with separate activation and readout lasers as is done when using mEOS [3]. The question of how fluorophore photophysics influences SMLM experiments is still under investigation [22], but this topic has recently been reviewed fairly comprehensively, taking into account the photoswitching characteristics of fluorescent proteins for SMLM [23].

We used mCitrine to perform SMLM of the growth factor ErbB3 in A431 epithelial carcinoma cells. A431 cells were chosen for this study in part because of their use in previous studies of the ErbB receptor system [24,25], and also because they tend to be very flat and form extended areas of membrane in contact with the coverslip, offering good conditions for SMLM. ErbB3 is a member of the epidermal growth factor receptor (EGFR) family, consisting of ErbB1 (EGFR), ErbB2 (also known as HER2), ErbB3, and ErbB4. The organization and dynamics of ErbB receptors is an important topic of study because overexpression and unrestrained activation of this family of receptors is implicated in cancer [26], including breast cancer [27]. Long thought to have no kinase activity, ErbB3 has recently been found to exhibit tyrosine kinase activity and to form homodimers and heterodimers with other ErbB receptors [28]. Such heterodimer formation between ErbB molecules can amplify signaling and appears to be an important feature of some cancer cells. In particular the ErbB2/ErbB3 heterodimer appears to be important for tumor cell proliferation in certain breast cancers [29]. High ErbB3 levels have been linked to resistance in cancer therapies which target ErbB1 or ErbB2 [30].

Given the importance of ErbB3 in cancer, an understanding of its organization and dynamics in the plasma membrane of tumor cells is critical. Super-resolution microscopy using single molecule localization reveals the coordinates of each ErbB3 receptor which is tagged with a YFP molecule. This data allows one to explore parameters such as clustering tendencies, an approach used successfully in studies of the T-cell receptor [31].

We have also included an additional single molecule super-resolution microscopy dataset acquired using the dye Alexa 532. This dye is more commonly used in (d)STORM studies [32] and is provided for purposes of comparison of the single molecule parameters. For this experiment we used an Alexa 532-labeled antibody to detect RNA molecules in the nucleus of a HeLa cell as previously described [33]. The raw data is useful in this context because it was acquired with the same microscope setup and detector. Compared to the YFP used in the other datasets, Alexa 532 has higher photon emission rates and exhibits less photobleaching.

The datasets have potential for extensive reuse. Complete raw data from SMLM experiments has typically not been published. The YFP data exhibits low signal to noise ratios, making data analysis a challenge. The data sets will be useful to investigators developing their own algorithms for SMLM, SOFI, and related methods. The data will also be useful for researchers investigating growth factor receptors such as ErbB3, as well as to those investigating other membrane proteins.

## Methods

### Cell lines and reagents

A431 cells (RRID: CVCL_0037) expressing mCitrine-ErbB3 and HeLa cells (RRID: CVCL_0030) were maintained in phenol red-free DMEM supplemented with 10 % FCS, 100 U/ml penicillin, 100 U/ml streptomycin, and L-glutamate (obtained from Invitrogen, Carlsbad, CA, USA) at 37 °C and 100% humidity. Mowiol 4-88 containing 1,4-diazabicyclo(2.2.2)octane (DABCO) was obtained from Fluka (St. Louis, MO, USA). Mercaptoethylamine (MEA) was obtained from Sigma (St. Louis, MO, USA).

### Sample preparation

Prior to SMLM experiments, A431 cells were grown on clean #1.5 coverslips for 12-18 hours. The cells were then washed with PBS, then fixed with 4% paraformaldehyde for 15 minutes at 4 °C. We then mounted the cells on clean slides using mowiol containing DABCO and 50-100 mM MEA, pH 8.5. Before microscopy, the mowiol was allowed to harden for 12-18 hours. The mowiol was freshly prepared according to standard procedures.

For labeling of transcription sites in the cell nucleus, HeLa cells were grown on #1.5 coverslips for 12-18 hours, then incubated for 5 minutes with 5-fluorouridine (Sigma) at a concentration of 10 μM. The cells were then fixed in 2% formaldehyde, permeabilized with 0.1% Triton X-100, and labeled using a mouse monoclonal anti-BrdU antibody (clone BU-33, Sigma). The anti-BrdU antibodies were then detected with a secondary anti-mouse antibody labeled with Alexa 532 (Invitrogen). The cells were mounted using freshly prepared mowiol containing DABCO and 50-100 mM MEA. Before microscopy, the mowiol was allowed to harden for 12-18 hours.

### Single molecule microscopy

For SMLM imaging, we used an IX71 microscope equipped with a planapochromatic 100×/1.35 NA oil immersion objective (Olympus, Tokyo, Japan) and a front-illuminated Ixon DU885 EMCCD camera under control of IQ software (Andor, Belfast, Northern Ireland) as previously described [34]. The excitation source was a 400 mW, 473 nm laser (Dragon laser, ChangChun, China), which was coupled to the microscope using a 0.39 NA multimode optical fiber. The fiber output was collimated using a 2 inch diameter, 60 mm FL lens (Thor Labs, Newton, New Jersey). The fiber was coupled into the microscope using an Olympus IX2-RFAL fluorescence illuminator, resulting in an evenly illuminated field. Fluorescence was observed using an Olympus U-MNIBA3 filter set (excitation 470 – 495 nm, dichroic 505 nm, emission 510 – 550 nm). In each experiment, a sequence of 1,419-10,000 images was acquired with an exposure time of 40 – 100 ms and an EM gain of 50-300. For imaging Alexa 532, we used a 1 W, 532 nm laser (Dragon laser) and an appropriate fluorescence emission filter (569-610 nm, Chroma) as previously described [33].

### Data analysis methods

We analyzed the data using ThunderSTORM [9,35] with the default settings. The default settings involve use of a wavelet-based filter for feature enhancement [36], followed by local maximum detection of single molecules in the filtered data. This is followed by fitting molecules in the raw data using a two-dimensional Gaussian function in integrated form [37] using maximum likelihood methods [38]. Gaussian functions have been found to be a good representation of the true PSF of a microscope [39]. For visualization of the results, we use an average shifted histogram approach [40]. If the camera calibration parameters (pixel size, photoelectrons per A/D count, base level, and EM gain) are correct, maximum likelihood fitting of an integrated Gaussian function will correctly return the number of photons detected from each molecule [9,37,38,41]. An integrated two dimensional Gaussian function can be written as

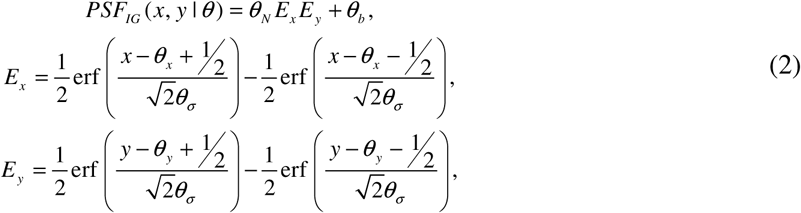

where *θ_x_*,*θ_y_* are the sub-pixel molecular coordinates, *θ*_*σ*_ is the standard deviation of the Gaussian function (i.e., the width), *θ_N_* is the total number of detected photons emitted by the molecule, and *θ_b_* is the background offset.

### Single molecule localization uncertainty

In ThunderSTORM the localization uncertainty is calculated for each detected molecule. This quantity can help one determine whether the molecule was well localized and whether it should be included in the final result. Let 
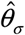
 be the standard deviation of a Gaussian function fitted to an imaged PSF in nm, *a* is the back-projected pixel size in nm (camera pixel size divided by system magnification), 
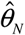
 is the estimate of the number of photons detected for a given molecule, and 
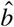
 is the background signal level in photons calculated as the standard deviation of the residuals between the raw data and the fitted PSF model. The uncertainty of estimates determined by maximum likelihood methods for the lateral position of a molecule is given by

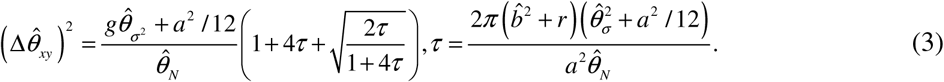

This formula is a modified form of the Thompson-Larson-Webb equation [11], and was derived by Rieger and Stallinga [42]. Finally, compensation for camera readout noise *r* and EM gain *g* was added following Quan, Zeng, and Huang [43], who suggested that when using EMCCD cameras, the correction factors should be set to *r* = 0, *g* = 2, and when using CCD or sCMOS cameras the correction factors should be set to *r* = *g* = 2.

### Super-resolution optical fluctuation imaging

Super-resolution optical fluctuation imaging (SOFI) is based on calculation of spatio-temporal cumulants over the input sequence of camera frames [44]. Assuming a non-fluctuating background and Gaussian additive noise, the n-th order cumulant (for *n* ≥ 2 and a time lag *τ*) can be written as

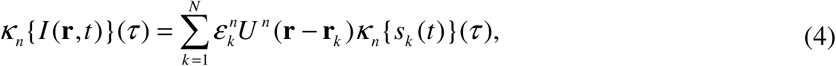

where *I* (**r**,*t*) is the detected intensity at position **r** and time *t*, *ε*_*k*_ is the molecular brightness of k-the emitter, *U^n^* (**r** − **r**_*k*_) is the PSF at the position **r**_*k*_, and *s*_*k*_ (*t*) denotes a normalized fluctuation sequence *s*_*k*_ (*t*)∈{0,1}. The PSF is raised to the n-th power, resulting in resolution increased by a factor of 
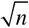
. After reweighting in frequency space, a resolution enhancement factor of *n* can be achieved [45], scaling linearly with the cumulant order. SOFI can be applied to any image sequence of stochastically blinking emitters acquired from a conventional widefield microscope if the emitters switch between at least two optically distinguishable states (a dark state and a bright state) and if sampling of the PSF fulfills the Nyquist–Shannon sampling theorem [46]. In comparison to STORM, SOFI tolerates higher densities of emitters and higher blinking rates [47], resulting in improved temporal resolution [48]. SOFI can be applied to the same datasets as SMLM analysis [47,49] offering an interesting complement to SMLM methods. Due to the entirely different image processing methods used, SOFI and SMLM are prone to different artifacts. Applying both processing methods to the same dataset reveals more information about the true structure and properties of the underlying sample. By combining multiple orders of the SOFI analysis, molecular parameters like molecular density, brightness, and on-time ratio can be extracted using the balanced SOFI method (bSOFI) [50]. The on-time ratio *ρ*_*on*_ describes the blinking rate of the fluorescent label. Assuming a two state blinking model where the emitter fluctuates between a bright state and a dark state, the on-time ratio is given as [38]

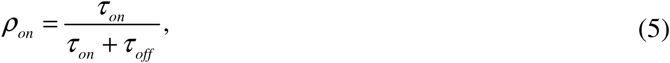

where the *τ_on_* and *τ_off_* are the characteristic lifetimes of the bright state and the dark state, respectively.

SOFI analysis was carried out as reported previously [49]. We used a custom written algorithm (Matlab, The Mathworks) based on the code of our SOFI simulation tool [51] and the bSOFI algorithm [50]. The sequence of camera frames was divided into subsequences of 500 frames each. The subsequences were processed separately in order to minimize the influence of photobleaching and the resulting SOFI images were averaged. Details about photobleaching correction for SOFI have recently been published [52]. SOFI relies on calculating higher order cumulants as described in the previous section. Calculating cumulants raises the molecular brightness to the n-th power (Eq. 3). SOFI’s non-linear response to brightness becomes an issue for cumulants of higher than second order where fluorescent spots of high brightness may mask less bright details. The balanced SOFI (bSOFI) algorithm linearizes the response to brightness [50] or to the detected intensity [49]. Throughout this work, the “n-th order bSOFI image” refers to an image calculated using the n-th order cumulant and applying the subsequent linearization according to the procedure described in [49].

## Super-resolution images

Figure 1 shows images of an A431 cell expressing mCitrine-ErbB3 (YFP dataset 1 [53]). Conventional widefield (WF, Fig. 1A), and SMLM (Fig. 1B) results are shown. Fig. 1C shows a color-coded density map, calculated by the bSOFI algorithm. This unique information cannot be obtained by conventional fluorescence microscopy. Fig. 1D shows the 4^th^ order bSOFI image.

**Fig. 1.**
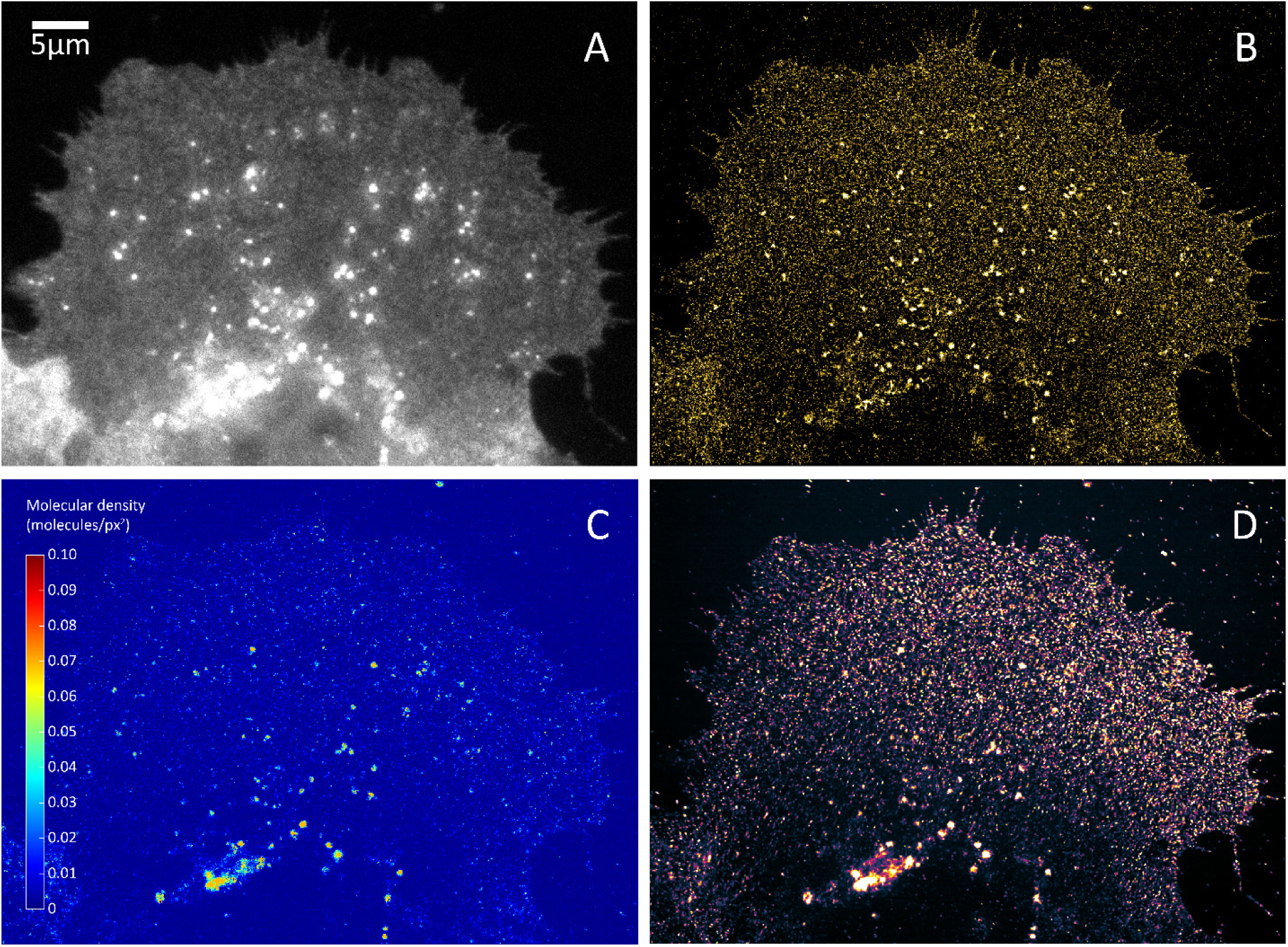
Super-resolution imaging of mCitrine-ErbB3 in A431 cells. (A) Conventional widefield. (B) SMLM. (C) Molecular density map. (D) 4^th^ order bSOFI.

Figure 2A shows a histogram of the number of photons detected from each YFP molecule (“intensity” in ThunderSTORM) for the cell shown in Fig 1. Fig. 2B shows a histogram of the localization uncertainty determined for each molecule for the cell shown in Fig 1. The localization uncertainty was calculated using Eq. 3. The two histograms were calculated using the *plot histogram* command in ThunderSTORM.

**Fig. 2.**
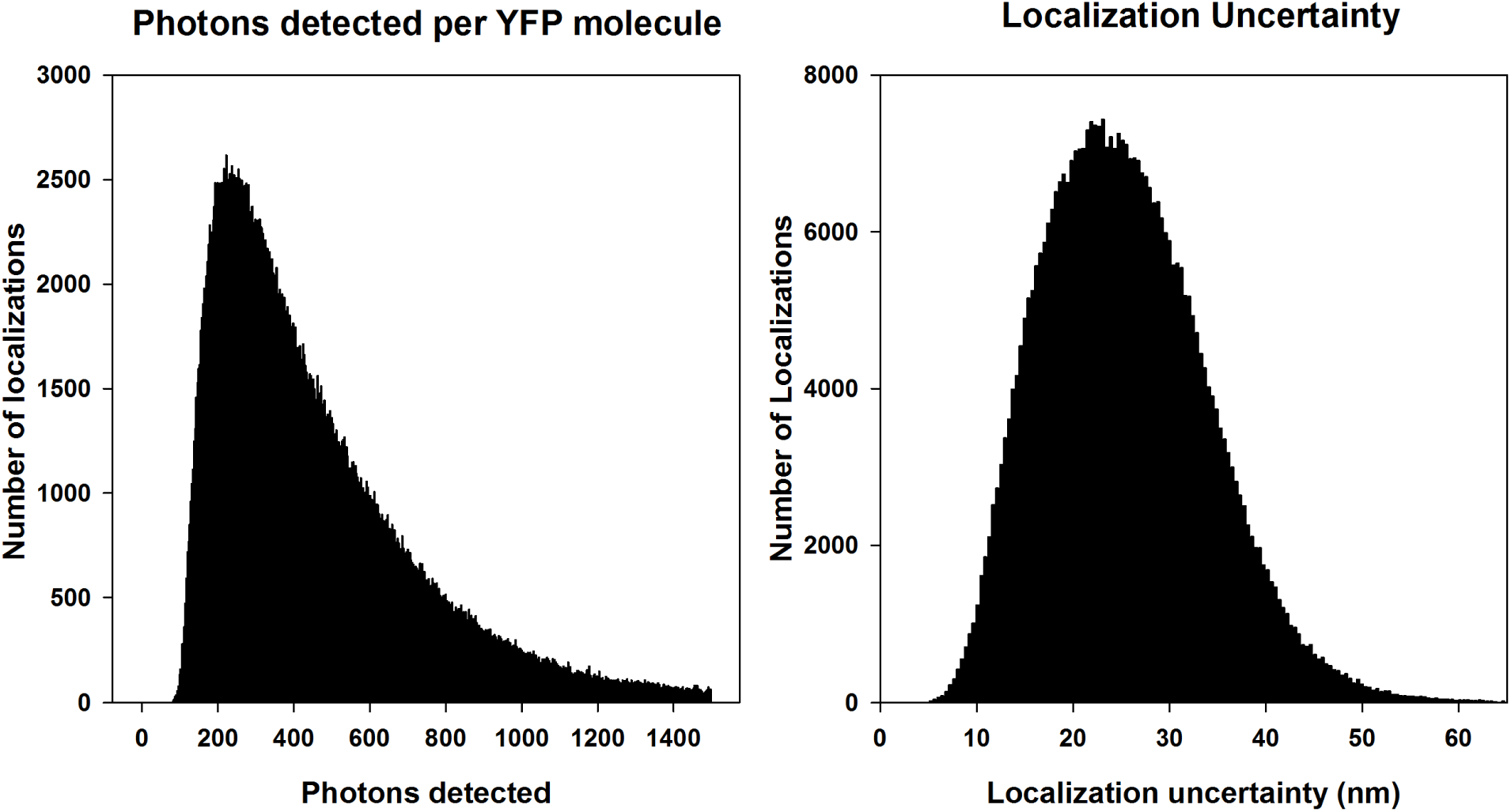
Quantification of molecular parameters from the experiment shown in Fig. 1. (A) Histogram of the number of photons detected from each YFP molecule. (B) Histogram of the localization uncertainty calculated for each YFP molecule.

Table 1 shows a list of quantitative parameters for the first 10 detected molecules as reported by ThunderSTORM for the experiment shown in Figure 1. Sigma (nm) is the standard deviation of the two-dimensional integrated Gaussian function fitted to the molecule, intensity (photons) is the number of photons detected from the molecule, offset (photons) is the background offset, SD of background (photons) is the standard deviation of the background, and localization uncertainty (nm) is the result of Equation 2 for each molecule. Recall that the full width at half max (FWHM) of a Gaussian function is related to its standard deviation by FWHM=2.35σ. The variation in parameters between molecules is usually attributed to differences in the local environment of each molecule such as oxygen concentration, and to factors such as the fluorophore orientation.

**Table 1.** Quantitative parameters for the first 10 detected molecules as reported by ThunderSTORM for the experiment shown in Fig. 1.

| Molecule number | Camera frame | x (nm) | y (nm) | sigma (nm) | intensity (photons) | offset (photons) | SD of Background (photons) | localization uncertainty (nm) |
| --- | --- | --- | --- | --- | --- | --- | --- | --- |
| 1 | 1 | 3743.17 | 28005.63 | 81.53 | 942 | 108 | 25 | 17.41 |
| 2 | 1 | 3880.95 | 31519.89 | 155.09 | 2014 | 68 | 21 | 23.33 |
| 3 | 1 | 4150.78 | 32662.21 | 60.03 | 433 | 81 | 21 | 17.75 |
| 4 | 1 | 4289.06 | 28407.32 | 36.90 | 407 | 155 | 32 | 12.97 |
| 5 | 1 | 4310.28 | 28737.99 | 103.00 | 1567 | 142 | 34 | 21.61 |
| 6 | 1 | 4615.06 | 23832.74 | 89.18 | 1186 | 73 | 22 | 14.60 |
| 7 | 1 | 4695.34 | 30060.77 | 102.05 | 1266 | 122 | 28 | 22.17 |
| 8 | 1 | 4812.40 | 30994.57 | 101.18 | 1051 | 115 | 24 | 22.61 |
| 9 | 1 | 4827.01 | 25960.59 | 83.02 | 717 | 80 | 20 | 19.35 |
| 10 | 1 | 5037.67 | 28686.08 | 149.32 | 2293 | 121 | 33 | 29.85 |

Figure 3 shows WF imaging of an A431 cell (Fig. 3A), along with identification of single molecules by ThunderSTORM (Fig. 3B, indicated by red dots), and the reconstructed SMLM result (Fig. 3C) (YFP dataset 2, [53]).

**Fig. 3.**
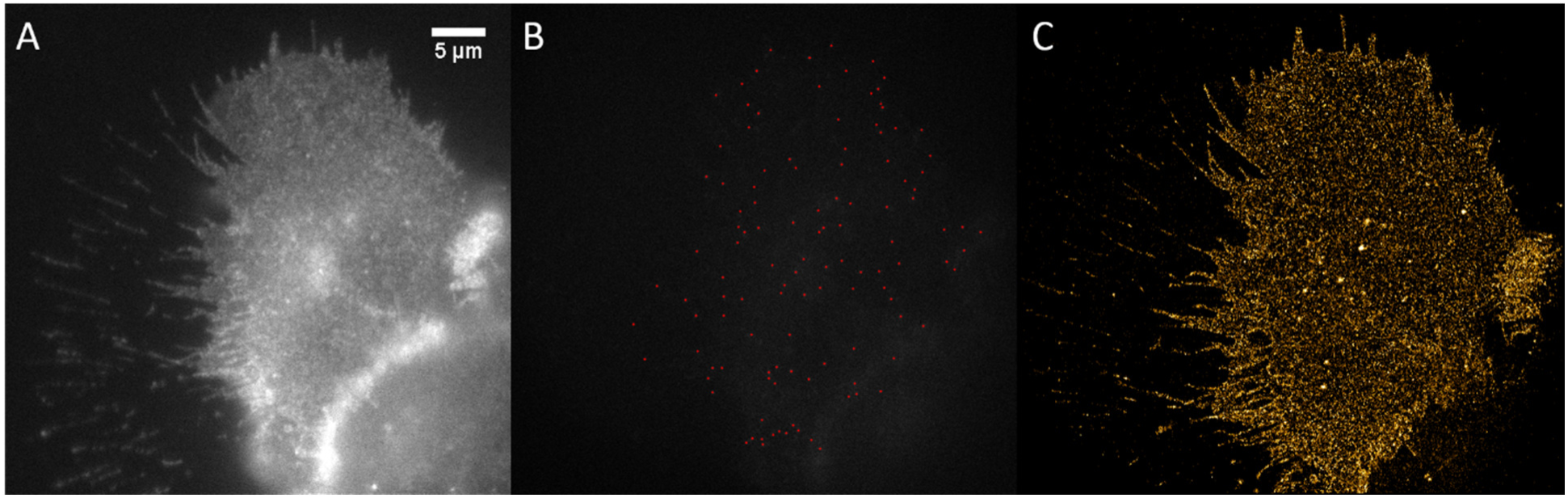
Super-resolution imaging of mCitrine-ErbB3 in A431 cells. (A) Conventional widefield. (B) Single frame of SMLM with detections indicated with red dots. (C) SMLM reconstruction.

Figure 4 shows WF imaging (Fig. 4A), and the reconstructed SMLM result (Fig. 4B) (YFP dataset 3, [53]). Figure 5 shows SOFI analysis for the cell shown in Figure 4. Second, third, and fourth order bSOFI images are shown in Fig. 5A-5C, as well as a density map (Fig 5D), photobleaching profile (Fig. 5E), and molecular on-time ratio (Fig 5F), where second, third, and fourth denotes the order of the cumulant used during the calculation of the bSOFI image. With increasing cumulant order of the SOFI analysis, spatial resolution generally increases, but the signal-to-background ratio (SBR) limits the spatial resolution achievable in practice. The situation is shown in detail in Fig 6B–D and in the line profiles in Fig 6F–H. The fourth order bSOFI image (Fig 6D) has higher spatial resolution compared to the second and third order bSOFI images (Fig 6B–C). The dashed lines in Fig 6F show the average value of the background of the bSOFI images, which increases for increasing order of SOFI analysis. In other words, increasing the cumulant order leads to a decrease in SBR which hampers the resolution enhancement. Note that we calculate linearized SOFI as previously described [49,50]. In the case of a relatively low density of emitters (Fig 6 F, H), SMLM achieved better spatial resolution. On the other hand, in Fig 6G, the SMLM analysis does not agree with the result from SOFI, suggesting that the local density of emitters was too high for successful single molecule identification and fitting in that particular location of the cell membrane.

Comparing the density maps in Fig 1D and Fig 5D, the sample in Fig. 5D exhibits an average density approximately 1.8 fold higher. The presence of more emitters in the sample (Fig 5D) leads to higher brightness which is likely the reason why the bSOFI image reconstruction was still successful despite the lower number of input frames.

**Fig. 4.**
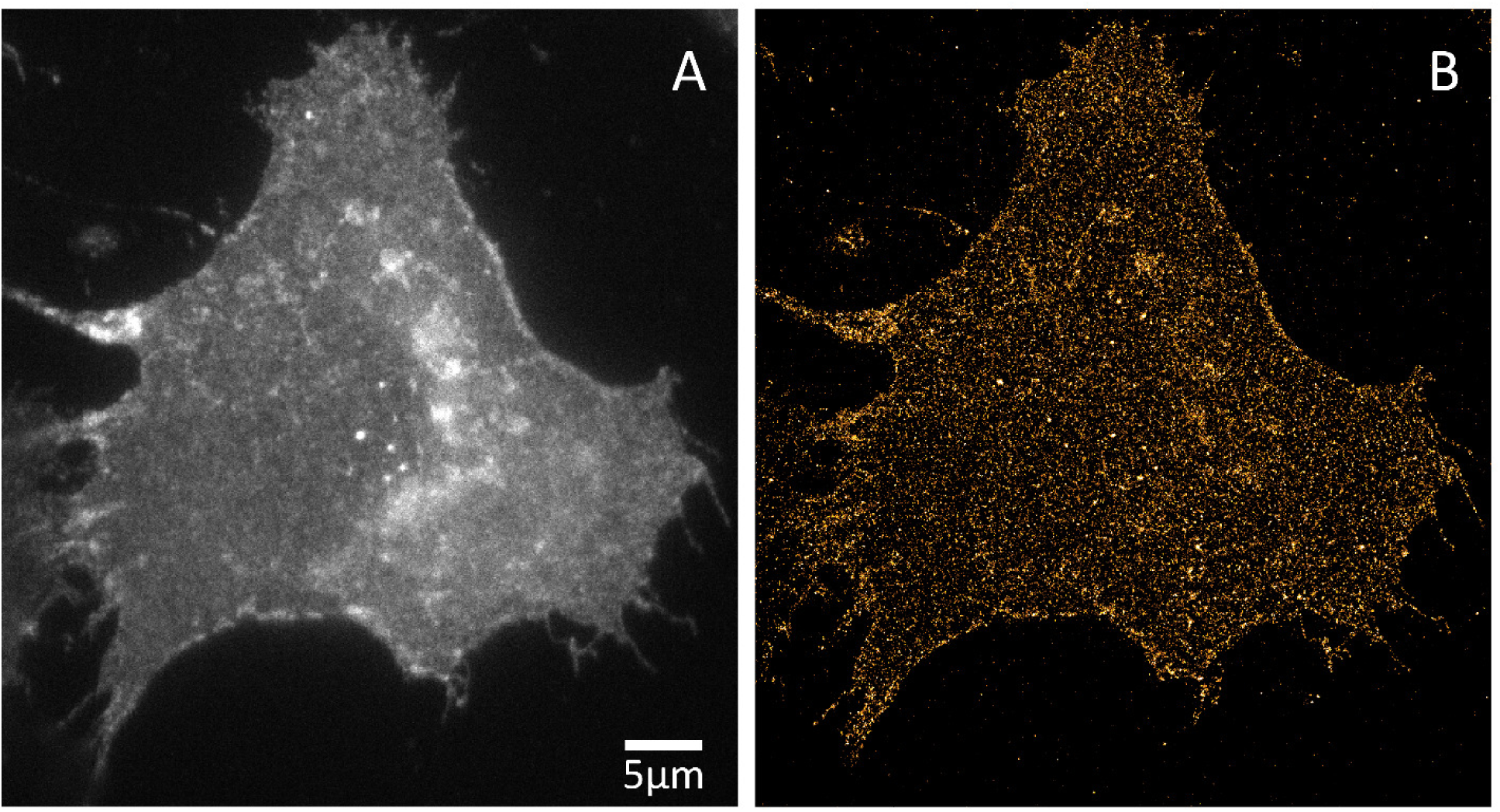
Super-resolution imaging of mCitrine-ErbB3 in A431 cells. (A) Conventional widefield. (B) SMLM reconstruction.

**Fig 5.**
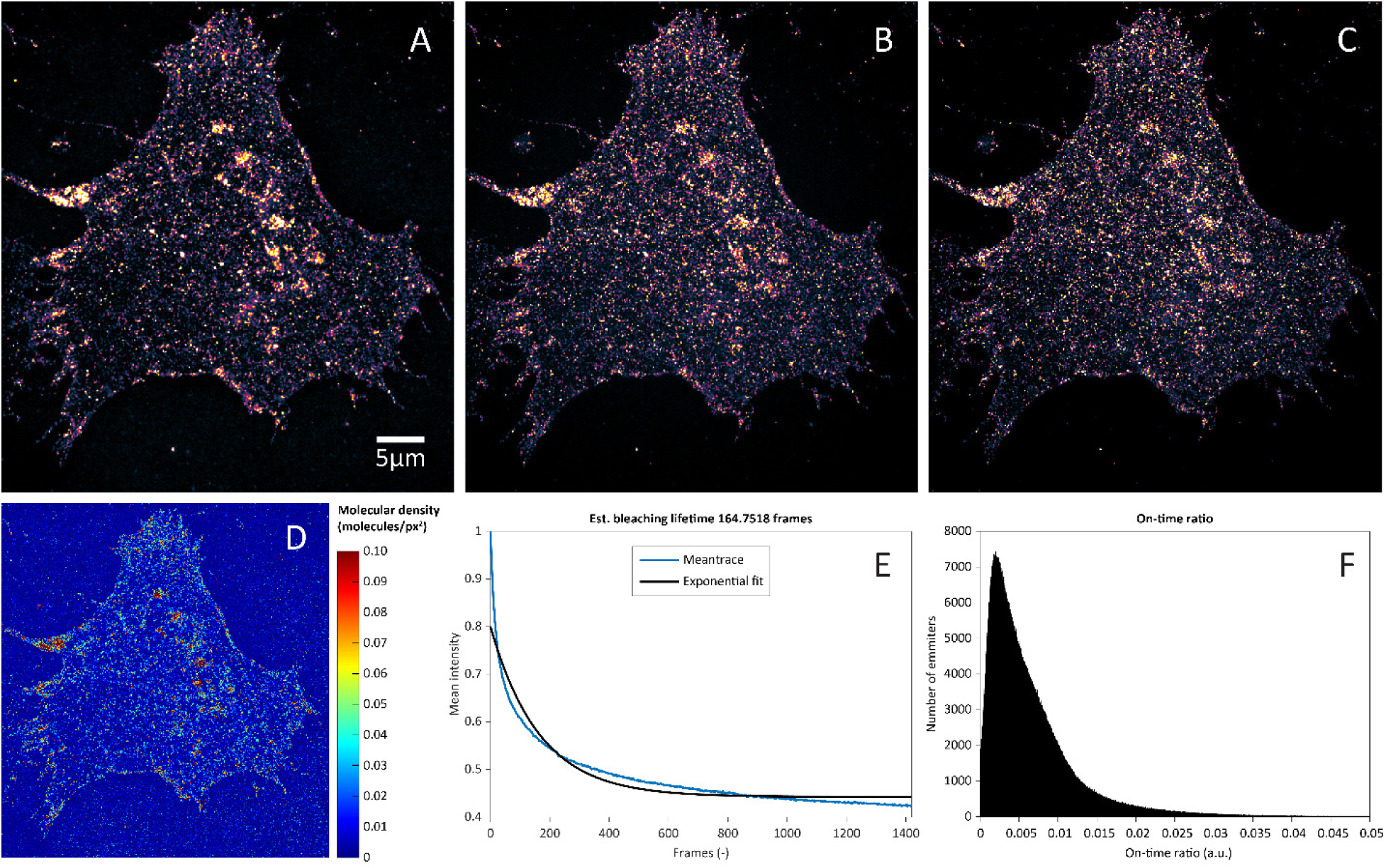
Super-resolution imaging of mCitrine-ErbB3 in A431 cells. (A,) (B) and (C) are second, third and fourth order bSOFI reconstruction, respectively. (D) Molecular density map estimated using bSOFI (E) Mean intensity trace of the raw image sequence (blue) with the exponential fit (black) used for photobleaching correction. (F) Histogram of the on-time ratio estimated using bSOFI algorithm.

**Fig 6.**
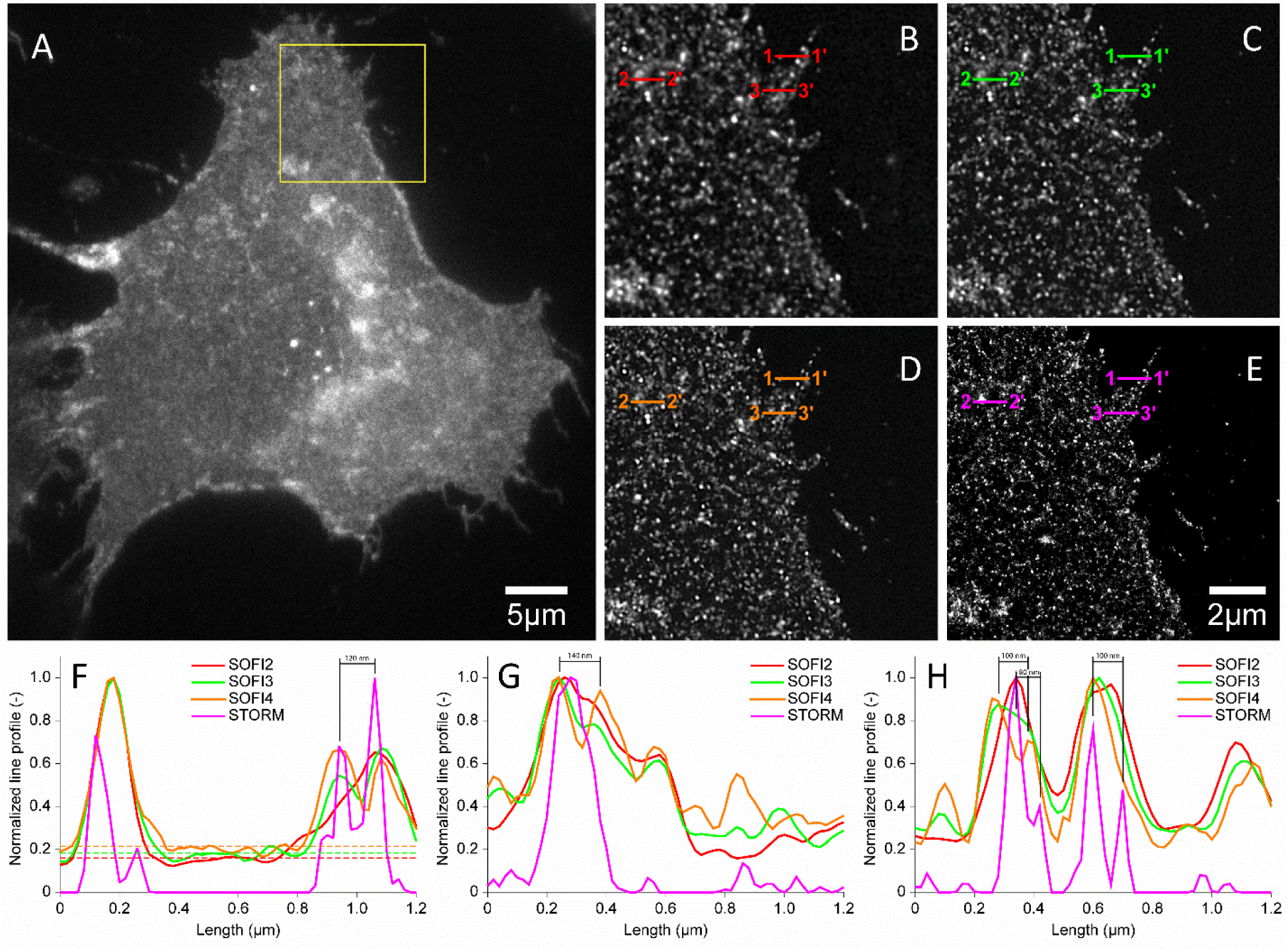
Super-resolution imaging of mCitrine-ErbB3 in A431 cells. (A) Conventional widefield. Region of interest marked in A by the yellow square processed by 2^nd^ order bSOFI (B), 3^rd^ order bSOFI (C), 4^th^ order bSOFI (D), SMLM (E). (F-H) Line profiles along the cuts 1-1’, 2-2’ and 3-3’ which correspond to examples of low density of emitters (F), high density (G), and medium density (H), respectively. Dashed lines in (F) represent the average value of the background of bSOFI images.

Table 2 shows a summary of the imaging conditions and quantitative parameters for the YFP and Alexa 532 datasets. Also shown are the relevant camera settings. The camera setting information should be entered into ThunderSTORM’s camera setup tab to ensure correct results.

**Table 2.** Summary of imaging conditions and quantitative parameters for the SMLM datasets.

| Data | Exp. time, ms | Pixel size, nm | e<sup>-</sup> per A/D count<sup>1</sup> | Base level, A/D counts | EM Gain | Frames | Total number of detections | Sigma, nm mean+/-SD | Loc. uncertainty, nm mean+/-SD |
| --- | --- | --- | --- | --- | --- | --- | --- | --- | --- |
| YFP data 1 (Fig. 1) | 50 | 80 | 3.6 | 414 | 150 | 10,000 | 482,778 | 86.9+/-29.0 | 25.6+/-8.6 |
| YFP data 2 (Fig. 2) | 100 | 80 | 3.6 | 414 | 50 | 6,366 | 224,175 | 84.1+/-26.3 | 27.3+/-8.7 |
| YFP data 3 (Figs. 4, 5, 6) | 50 | 80 | 3.6 | 414 | 150 | 1,419 | 452,498 | 84.2+/-24.3 | 28.9+/-8.1 |
| YFP data 4 | 100 | 80 | 3.6 | 414 | 100 | 3,922 | 159,463 | 81.6+/-24.1 | 25.9+/-8.0 |
| Alexa 532 data | 30 | 80 | 1.5 | 396 | 50 | 20,000 | 1,128,322 | 121.5+/-47.6 | 20.6+/-7.5 |

<sup>1</sup>photoelectrons per analog to digital converter count

## Re-use potential

Super-resolution microscopy algorithms are under active development [10]. Researchers engaged in algorithm development may use this dataset to help develop and fine tune their methods. Since the true positions of the molecules remain unknown, the results from ThunderSTORM may be taken as the reference data for comparison purposes. ThunderSTORM offers an analysis tool which compares reference data and experimental data and computes several quantities which can be used to quantitatively evaluate algorithm performance. A detailed example of use is provided in the supplementary information.

## Availability of source code and requirements

Project name: ThunderSTORM v1.3

Project home page: <u>http://zitmen.github.io/thunderstorm/</u>

Operating system: platform independent

Programming language: Java

Other requirements: Image J <u>https://imagej.nih.gov/ij/</u>

License: GNU General Public License v3.0

## Availability of data

All raw and analyzed data are available in the GigaScience repository, GigaDB [53].

## Abbreviations

(d)STORM: (direct) stochastic optical reconstruction microscopy
FWHM: full width at half maximum
GFP: green fluorescent protein
NA: numerical aperture
PALM: photoactivated localization microscopy
PSF: point spread function
SMLM: single molecule localization microscopy
SOFI: stochastic optical fluctuation imaging
WF: wide field
YFP: yellow fluorescent protein

## Author Contributions

TL: analyzed data, developed computer code, wrote the paper

JP: analyzed data, developed computer code

KF: supervised research

TL: supervised research

GH: conceived project, acquired data, analyzed data, supervised research, wrote the paper

## Acknowledgements

Epithelial carcinoma A431 cells expressing mCitrine-ErbB3 were a kind gift from Dr. Donna Arndt-Jovin and Dr. Tom Jovin of the Max Planck Institute for Biophysical Chemistry (Göttingen, Germany). We thank Peter W. Winter for assistance with microscopy, and Pavel Křížek, Josef Borkovec, Zdeněk Švindrych, and Martin Ovesný for assistance with microscopy, data analysis, and programming. We thank Evgeny Smirnov for assistance with sample preparation.

## Supplementary information: Example of data re-use: quantitative evaluation of SMLM algorithms using ThunderSTORM

### 1.1 Introduction

ThunderSTORM [1] can compare the results of two SMLM analysis methods by matching, within a user-selected tolerance, each localization in a results table with the nearest corresponding molecule in a second results table. In this way a new algorithm can be tested against a standard method. Typically this is done with simulated data, but this approach can be used with experimental data as well. The localization microscopy datasets presented here can be used to develop, evaluate, and validate new SMLM algorithms. When working with simulated data, the true positions of the simulated molecules are known (ground-truth positions). When working with experimental data we must use the results from a standard algorithm as the reference data and test this against the results of an algorithm under development. In ThunderSTORM, the reference dataset is always called the “ground-truth” data.

### 1.2 Counting localized and missed molecules

The process of performance evaluation starts by pairing the localized molecules with the closest molecule in the ground-truth data (or the reference data set), see Figure 1. The numbers of correctly and incorrectly identified molecules are counted as follows. If the distance between the paired molecules is smaller than a user-specified radius, then the localization is counted as a true positive (TP) detection and the localized molecule is associated with the ground-truth position. If the distance is greater than or equal to that radius, then the localization is counted as a false positive (FP) detection. Ground-truth molecules which were not associated with the localized molecules are counted as false negatives (FNs). With a growing density of molecules it becomes more important how the algorithm performs the matching. To solve the problem of finding the correct matching between localized molecules and the ground-truth data, we use the Gale-Shapley algorithm [2].

### 1.3. Precision and recall

Statistical measures related to the number of correctly or incorrectly detected molecules, or missed molecules, are the recall (r) (also called sensitivity) and the precision (p) (also called positive predictive value) [3–5]. Their definitions are given by

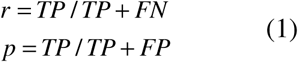

Recall measures the fraction of correctly identified molecules, and precision measures the portion of correctly identified molecules out of the set of all localizations. The theoretical optimum is achieved for values of recall and precision both equal to 1.0.

### 1.4 F1 score

For purposes of comparison between multiple algorithms, it is convenient to combine precision and recall into a single measure of performance with some trade-offs between both values. A traditional method for this applies the F1 score [3,4], defined as

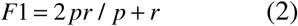

Values of the F1 score close to zero indicate both bad recall and precision while values approaching 1.0 signify a good ratio between recall and precision.

### 1.5 Jaccard index

Another measure suitable for comparing similarity and diversity of sets of samples is the Jaccard index defined by the formula

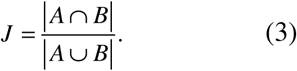

Here A is the set of ground-truth molecular positions, B is the set of all molecular positions localized by processing the data, intersection ∣*A ∩ B*∣ = *TP* gives the number of true positive detections, union ∣*A ∪ B*∣ = *TP* + *FP* + *FN*. The Jaccard index ranges from zero to one and a theoretical optimum is achieved for values of the Jaccard index equal to 1.0.

### 1.6 RMS distance

For all molecules identified as true positives, we also calculate the root-mean square distance between the ground-truth positions of the molecules and their localizations.

### 1.7 Performance Evaluation

If ground truth molecular positions are available, ThunderSTORM can evaluate the performance of different processing methods. Ground truth data can be created by ThunderSTORM’s generator of simulated data, or imported from a file by selecting `Plugins → ThunderSTORM → Import/Export → Import ground-truth.` To start evaluation, select `Plugins → ThunderSTORM → Performance testing → Performance evaluation`, enter the tolerance radius for accepting true-positive detections, and click OK. The results indicate true-positive detections (green), false positive detections (red), false negatives (orange), and related statistical measures (e.g., recall, precision, F1 score).

**Fig S1.**
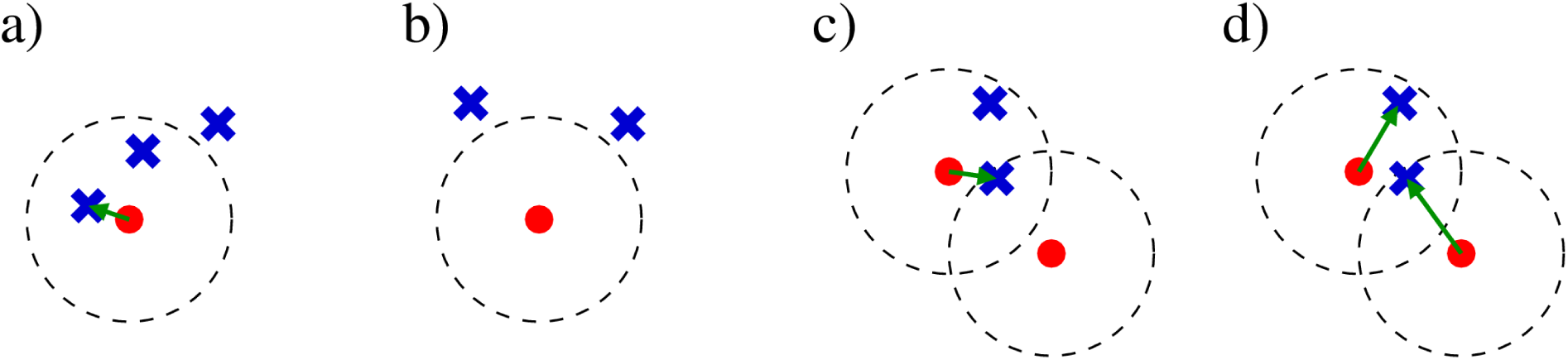
Counting localized and missed molecules. Red dot - ground-truth position of a molecule, blue cross - localized molecule, green arrow - association of a localized molecule with ground-truth position, dashed circle - detection tolerance radius. a) 1 TP + 1 FP, b) 1 FN + 2 FP, c-d) example of a situation, where c) greedy approach fails by finding 1 TP + 1 FP + 1 FN, and where d) Gale-Shapley algorithm finds a correct solution with 2 TP.

**Fig S2.**
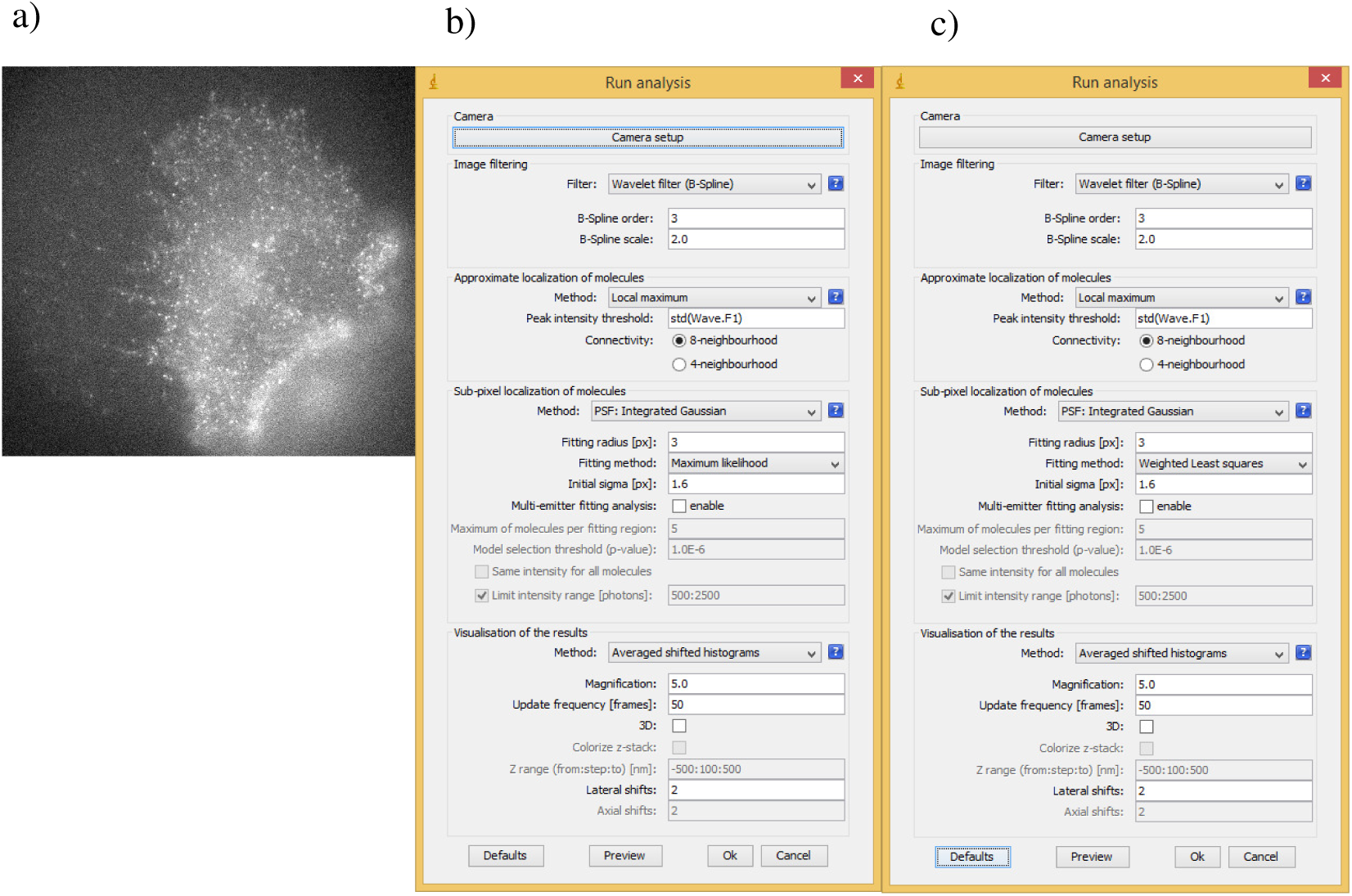
a) input image (frame 100 of ‘YFP dataset 2’), b) ThunderSTORM setup, default settings with maximum likelihood fitting method selected, c) ThunderSTORM setup, default settings with weighted least squares fitting method selected.

**Fig S3.**
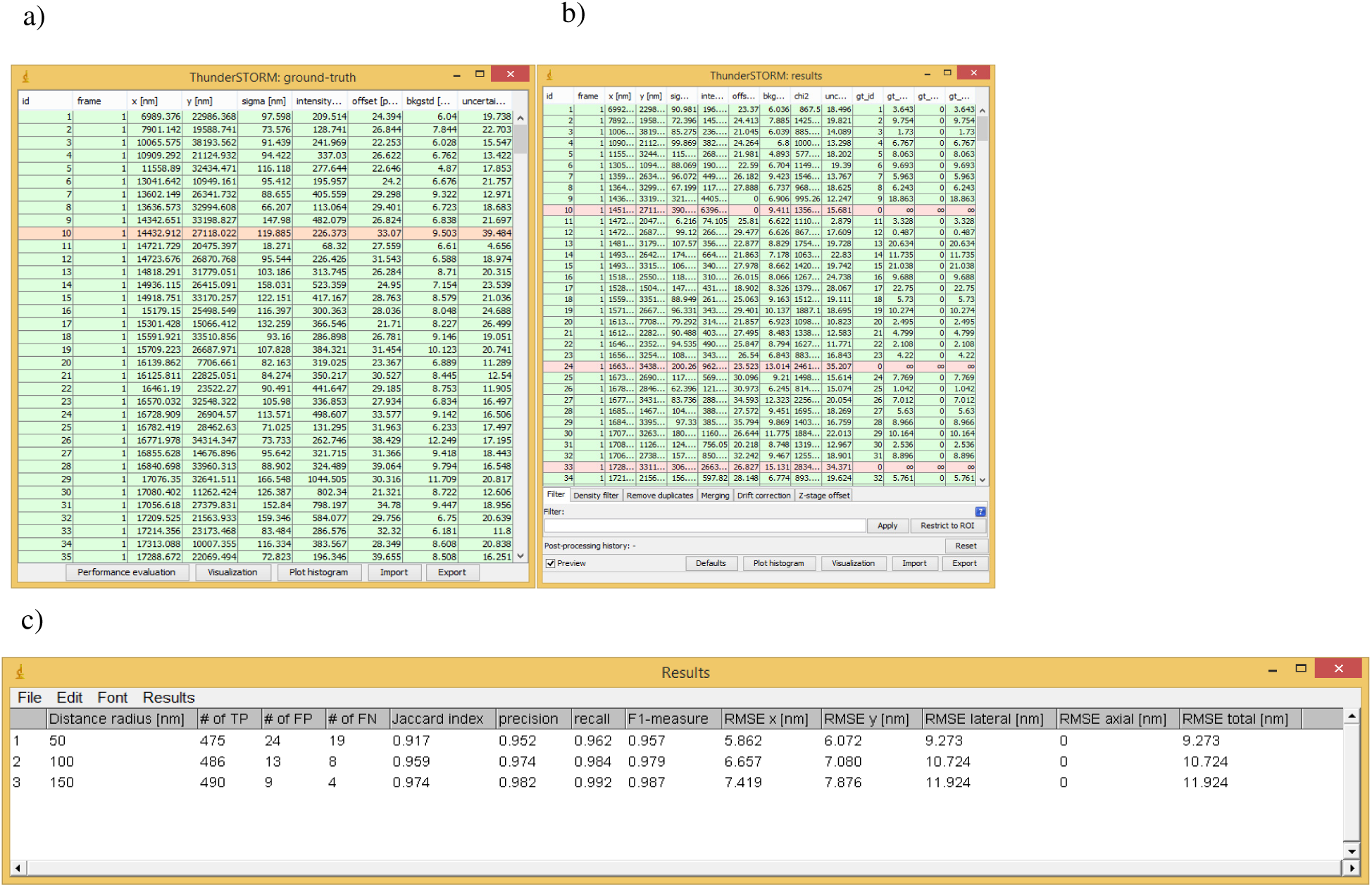
a) ThunderSTORM results table using maximum likelihood fitting. b) ThunderSTORM results table using weighted least squares fitting. The results indicate true-positive detections (green), false positive detections (red), and false negatives (orange). c) Table of results when varying the molecule matching tolerance. Statistics are calculated which quantitatively compares the two results tables.

## References

[1] Huang B, Bates M, Zhuang X. Super-resolution fluorescence microscopy. Annu. Rev. Biochem. 2009;78:993–1016.

[2] Hell SW, Sahl SJ, Bates M, Zhuang X, Heintzmann R, Booth MJ, et al. The 2015 super-resolution microscopy roadmap. J. Phys. D. Appl. Phys. 2015;48:443001.

[3] Betzig E, Patterson GH, Sougrat R, Lindwasser OW, Olenych S, Bonifacino JS, et al. Imaging intracellular fluorescent proteins at nanometer resolution. Science. 2006;313:1642–5.

[4] Wiedenmann J, Ivanchenko S, Oswald F, Schmitt F, Röcker C, Salih A, et al. EosFP, a fluorescent marker protein with UV-inducible green-to-red fluorescence conversion. Proc. Natl. Acad. Sci. U. S. A. 2004;101:15905–10.

[5] Rust MJ, Bates M, Zhuang X. Sub-diffraction-limit imaging by stochastic optical reconstruction microscopy (STORM). Nat. Methods. 2006;3:793–5.

[6] Huang B, Wang W, Bates M, Zhuang X. Three-dimensional super-resolution imaging by stochastic optical reconstruction microscopy. Science. 2008;319:810–3.

[7] Heilemann M, van de Linde S, Schüttpelz M, Kasper R, Seefeldt B, Mukherjee A, et al. Subdiffraction-resolution fluorescence imaging with conventional fluorescent probes. Angew. Chemie Int. Ed. 2008;47:6172–6.

[8] Dempsey GT, Vaughan JC, Chen KH, Bates M, Zhuang X. Evaluation of fluorophores for optimal performance in localization-based super-resolution imaging. Nat. Methods. 2011;8:1–14.

[9] Ovesný M, Křížek P, Borkovec J, Švindrych Z, Hagen GM. ThunderSTORM: A comprehensive ImageJ plug-in for PALM and STORM data analysis and super-resolution imaging. Bioinformatics. 2014;30.

[10] Sage D, Kirshner H, Pengo T, Stuurman N, Min J, Manley S, et al. Quantitative evaluation of software packages for single-molecule localization microscopy. Nat. Methods. 2015;12:717–24.

[11] Thompson RE, Larson DR, Webb WW. Precise nanometer localization analysis for individual fluorescent probes. Biophys. J. 2002;82:2775–2783.

[12] Fox-Roberts P, Marsh R, Pfisterer K, Jayo A, Parsons M, Cox S. Local dimensionality determines imaging speed in localization microscopy. Nat. Commun. 2017;8:13558.

[13] Dickson RM, Cubitt AB, Tsien RY, Moerner WE. On/off blinking and switching behaviour of single molecules of green fluorescent protein. Nature. 1997;388:355–358.

[14] Lemmer P, Gunkel M, Baddeley D, Kaufmann R, Urich A, Weiland Y, et al. SPDM: light microscopy with single-molecule resolution at the nanoscale. Appl. Phys. B Lasers Opt. 2008;93:1–12.

[15] Lemmer P, Gunkel M, Weiland Y, Muller P, Baddeley D, Kaufmann R, et al. Using conventional fluorescent markers for far-field fluorescence localization nanoscopy allows resolution in the 10-nm range. J. Microsc. 2009;235:163–71.

[16] Biteen JS, Thompson MA, Tselentis NK, Bowman GR, Shapiro L, Moerner WE. Super-resolution imaging in live Caulobacter crescentus cells using photoswitchable EYFP. Nat. Methods. 2008;5:947–9.

[17] Lew MD, Lee SF, Ptacin JL, Lee MK, Twieg RJ, Shapiro L, et al. Three-dimensional superresolution colocalization of intracellular protein superstructures and the cell surface in live Caulobacter crescentus. Proc. Natl. Acad. Sci. U. S. A. 2011;108:E1102–10.

[18] Jusuk I, Vietz C, Raab M, Dammeyer T, Tinnefeld P., Super-Resolution Imaging Conditions for enhanced Yellow Fluorescent Protein (eYFP) Demonstrated on DNA Origami Nanorulers. Sci. Rep. 2015;5:14075.

[19] Křížek P, Raška I, Hagen GM. Minimizing detection errors in single molecule localization microscopy. Opt. Express. 2011;19:3226–35.

[20] Kaufmann R, Piontek J, Grüll F, Kirchgessner M, Rossa J, Wolburg H, et al. Visualization and quantitative analysis of reconstituted tight junctions using localization microscopy. PLoS One. 2012;7:e31128.

[21] Griesbeck O, Baird GS, Campbell RE, Zacharias DA, Tsien RY. Reducing the environmental sensitivity of yellow fluorescent protein. J. Biol. Chem. 2001;276:29188–29194.

[22] Pennacchietti F, Gould TJ, Hess ST. The role of probe photophysics in localization-based superresolution microscopy. Biophys. J. 2017;113:2037–54.

[23] Shcherbakova DM, Sengupta P, Lippincott-Schwartz J, Verkhusha V V. Photocontrollable fluorescent proteins for superresolution imaging. Annu. Rev. Biophys. 2014;43:303–29.

[24] Nagy P, Arndt-Jovin DJ, Jovin TM., Small interfering RNAs suppress the expression of endogenous and GFP-fused epidermal growth factor receptor (erbB1) and induce apoptosis in erbB1-overexpressing cells. Exp. Cell Res. 2003;285:39–49.

[25] Hagen GM, Caarls W, Lidke KA, De Vries AHB, Fritsch C, Barisas BG, et al. Fluorescence recovery after photobleaching and photoconversion in multiple arbitrary regions of interest using a programmable array microscope. Microsc. Res. Tech. 2009;72:431–40.

[26] Yarden Y, Sliwkowski. Untangling the ErbB signaling network. Nat. Rev. Mol. Cell Biol. 2001;2:127–37.

[27] Naidu R, Yadav M, Nair S, Kutty MK. Expression of c-erbB3 protein in primary breast carcinomas. Br. J. Cancer. 1998;78:1385–90.

[28] Steinkamp MP, Low-Nam ST, Yang S, Lidke KA, Lidke DS, Wilson BS. ErbB3 is an active tyrosine kinase capable of homo‐ and heterointeractions. Mol. Cell. Biol. 2014;34:965–77.

[29] Holbro T, Beerli RR, Maurer F, Koziczak M, Barbas CF, Hynes NE., The ErbB2/ErbB3 heterodimer functions as an oncogenic unit: ErbB2 requires ErbB3 to drive breast tumor cell proliferation. Proc. Natl. Acad. Sci. U. S. A. 2003;100:8933–8938.

[30] Sithanandam G, Anderson LM., The ErbB3 receptor in cancer and cancer gene therapy. Cancer Gene Ther. 2008;15:413–48.

[31] Williamson DJ, Owen DM, Rossy J, Magenau A, Wehrmann M, Gooding JJ, et al. Pre-existing clusters of the adaptor Lat do not participate in early T cell signaling events. Nat Immunol. 2011;12:655–62.

[32] Heilemann M, Linde S van de, Mukherjee A, Sauer M. Super-resolution imaging with small organic fluorophores. Angew. Chemie Int. Ed. 2009;48:6903–6908.

[33] Smirnov E, Borkovec J, Kováčik L, Svidenská S, Schröfel A, Skalníková M, et al. Separation of replication and transcription domains in nucleoli. J. Struct. Biol. 2014;188:259–66.

[34] Křížek P, Raška I, Hagen GM. Flexible structured illumination microscope with a programmable illumination array. Opt. Express. 2012;20:24585–99.

[35] Ovesný M, Křížek P, Borkovec J, Švindrych Z, Hagen GM. Image analysis for single-molecule localization microscopy. In: Diaspro A, van Zandvoort Marc A. M. J., editors. Super-Resolution Imaging Biomed., Boca Raton, Florida: CRC Press; 2016, p. 79–97.

[36] Izeddin I, Boulanger J, Racine V, Specht CG, Kechkar A, Nair D, et al. Wavelet analysis for single molecule localization microscopy. Opt. Express. 2012;20:2081–95.

[37] Huang F, Schwartz SL, Byars JM, Lidke KA. Simultaneous multiple-emitter fitting for single molecule super-resolution imaging. Biomed. Opt. Express. 2011;2:1377–93.

[38] Mortensen KI, Churchman LS, Spudich JA, Flyvbjerg H. Optimized localization analysis for single-molecule tracking and super-resolution microscopy. Nat. Methods. 2010;7:377–381.

[39] Stallinga S, Rieger B. Accuracy of the gaussian point spread function model in 2D localization microscopy. Opt. Express. 2010;18:24461–76.

[40] Scott DW. Averaged shifted histograms: effective nonparametric density estimators in several dimensions. Ann. Stat. 1985;13:1024–40.

[41] Smith CS, Joseph N, Rieger B, Lidke KA. Fast, single-molecule localization that achieves theoretically minimum uncertainty. Nat. Methods. 2010;7:373–375.

[42] Rieger B, Stallinga S. The lateral and axial localization uncertainty in super-resolution light microscopy. Chemphyschem. 2014;15:664–70.

[43] Quan T, Zeng S, Huang Z-L. Localization capability and limitation of electron-multiplying charge-coupled, scientific complementary metal-oxide semiconductor, and charge-coupled devices for superresolution imaging. J. Biomed. Opt. 2010;15:66005.

[44] Dertinger T, Colyer R, Iyer G, Weiss S, Enderlein J. Fast, background-free, 3D super-resolution optical fluctuation imaging (SOFI). Proc. Natl. Acad. Sci. U. S. A. 2009;106:22287–22292.

[45] Dertinger T, Colyer R, Vogel R, Enderlein J, Weiss S. Achieving increased resolution and more pixels with superresolution optical fluctuation imaging (SOFI). Opt. Express. 2010;18:18875–85.

[46] Heintzmann R. Band-limit and appropriate sampling in microscopy. In: Celis Julio E, editor. Cell Biol. A Lab. Handb., Elsevier Academic Press; 2006, p. 29–36.

[47] Geissbuehler S, Dellagiacoma C, Lasser T. Comparison between SOFI and STORM. Biomed. Opt. Express. 2011;2:408–20.

[48] Geissbuehler S, Sharipov A, Godinat A, Bocchio NL, Sandoz PA, Huss A, et al. Live-cell multiplane three-dimensional super-resolution optical fluctuation imaging. Nat. Commun. 2014;5:5830.

[49] Deschout H, Lukes T, Sharipov A, Szlag D, Feletti L, Vandenberg W, et al. Complementarity of PALM and SOFI for super-resolution live-cell imaging of focal adhesions. Nat. Commun. 2016;7:13693.

[50] Geissbuehler S, Bocchio NL, Dellagiacoma C, Berclaz C, Leutenegger M, Lasser T. Mapping molecular statistics with balanced super-resolution optical fluctuation imaging (bSOFI). Opt. Nanoscopy. 2012;1:4.

[51] Girsault A, Lukeš T, Sharipov A, Geissbuehler S, Leutenegger M, Vandenberg W, et al. SOFI simulation tool: A software package for simulating and testing super-resolution optical fluctuation imaging. PLoS One. 2016;11:e0161602.

[52] Peeters Y, Vandenberg W, Duwé S, Bouwens A, Lukeš T, Ruckebusch C, et al. Correcting for photodestruction in super-resolution optical fluctuation imaging. Sci. Rep. 2017;7:July 2017.

[53] Lukeš, T; Pospíšil, J; Fliegel, K; Lasser, T; Hagen, G M. Supporting data for “Quantitative super-resolution single molecule microscopy dataset of YFP-tagged growth factor receptors.” GigaScience Database. 2018. doi:http://dx.doi.org/10.5524/100400.

## Supplementary References

[1] Ovesný M, Křížek P, Borkovec J, Švindrych Z, Hagen GM. ThunderSTORM: A comprehensive ImageJ plug-in for PALM and STORM data analysis and super-resolution imaging. Bioinformatics 2014;30.

[2] Gale D, Shapley LS. College Admissions and the Stability of Marriage. Am Math Mon. 1962;69:9–15.

[3] Tan P-N, Steinbach M, Kumar V. Introduction to data mining. Pearson Addison Wesley; 2005.

[4] Křížek P, Raška I, Hagen GM. Minimizing detection errors in single molecule localization microscopy. Opt Express. 2011;19:3226–35.

[5] Wolter S, Löschberger A, Holm T, Aufmkolk S, Dabauvalle M-C, van de Linde S, et al. rapidSTORM: accurate, fast open-source software for localization microscopy. Nat Methods. 2012;9:1040–1.

